# From synaptic activity to human in vivo quantification of neurotransmitter dynamics: a neural modelling approach

**DOI:** 10.1101/2021.03.11.434540

**Authors:** Caroline A. Lea-Carnall, Wael El-Deredy, Stephen R. Williams, Charlotte J. Stagg, Nelson J. Trujillo-Barreto

## Abstract

Understanding the role of neurotransmitters glutamate and GABA during normal and abnormal brain function and under external stimulation in humans are critical neuroscientific and clinical goals. The recent development of functional 1H-Magnetic resonance spectroscopy (fMRS) has allowed us to study neurotransmitter activity *in vivo* for the first time. However, the physiological basis of the observed fMRS signal remains unclear. It has been proposed that fMRS detects shifts in metabolite concentrations as they move from presynaptic vesicles, where they are largely invisible, to extracellular and cytosolic pools, where they are visible.

Here we bridge the gap between neural dynamics and fMRS by developing a mean-field model to link the neurotransmitter dynamics at the microscopic-level to the macroscopic-level imaging measurements. GABA and glutamate are described as cycling between three metabolic pools: in the vesicles; active in the cleft; or undergoing recycling in the astrocytic or neuronal cytosol. We interrogate the model by applying a current to manipulate the mean membrane potential and firing rate of the neural populations.

We find that by disregarding the contribution from the vesicular pool, our model predicts activity-dependent changes in the MRS signal, which are consistent with reported empirical findings. Further, we show that current magnitude and direction has a selective effect on the GABA/glutamate-MRS signal: inhibitory stimulation leads to reduction of both metabolites, whereas excitatory stimulation leads to increased glutamate and decreased GABA. In doing so, we link neural dynamics and fMRS and provide a mechanistic account for the activity-dependent change in the observed MRS signal.

**Key Points Summary:**

1. The recent development of functional 1H-Magnetic resonance spectroscopy (fMRS) has allowed us to study neurotransmitter activity *in vivo* for the first time in humans. However, the physiological basis of the observed fMRS signal is unclear.
2. It has been proposed that fMRS detects shifts in metabolite concentrations as they move from presynaptic vesicles, where they are largely invisible to MRS, to extracellular and cytosolic pools, where they are visible to MRS.
3. We test this hypothesis using a mean field model which links the neural dynamics of neurotransmitters at the microscopic-level to the macroscopic-level imaging measurements obtained in experimental studies.
4. By disregarding activity in the vesicular pool, our model can generate activity-dependent changes in the MRS signal in response to stimulation which are consistent with experimental findings in the literature.
5. We provide a mechanistic account for the activity-dependent change in observed neurotransmitter concentrations using MRS.

## 1 Introduction

Understanding the physiological changes driving human behaviour and clinical conditions is of central importance to clinical and neuroscientific fields. However, the need for non-invasive methods to quantify neural activity in humans makes this question inherently challenging. MRS is the only technique that allows quantification of brain metabolites non-invasively, providing a method to directly measure the concentrations of the brain’s major neurotransmitters (NT) glutamate (Glu) and *γ*-aminobutyric acid (GABA) in humans [1]. However, it is not clear how to interpret the resulting signals in terms of the underlying cellular mechanisms. Here, we use mean-field theory to provide a macroscopic level account of the basis of Glu and GABA fMRS signal changes due to current stimulation. We then use the model to predict the precise metabolic pools that give rise to the in-vivo measurements.

Traditionally, MRS has been used to provide a static snapshot of neurotransmitter levels in the brain, predominantly used to allow comparison between healthy and diseased states. However, there is now increasing use of dynamic measurements to quantify metabolic responses to stimuli on much shorter timescales of seconds to minutes, so-called functional MRS (see [2, 3], for reviews). Following earlier work in the visual system, fMRS applications have now been expanded to a range of other paradigms including motor, cognitive, and pain stimuli, as reviewed by Jelen *et al.* and Mullins [2, 3]. In many of these studies, authors report changes in neurochemicals of up to 10-14% in response to short bouts of stimulation. In addition to studies focussed on healthy volunteers, the utility of fMRS techniques in clinical populations is also being increasingly explored. For example, a recent fMRS study in the anterior cingulate cortex (ACC) showed that the expected Glu increase in response to a Stroop task was significantly decreased in people with schizophrenia and major depressive disorder [4]. Given that the ACC has been substantially implicated in schizophrenia, with increased Glu and glutamine (Gln) levels observed commonly, fMRS is therefore able to significantly aid our understanding of the pathophysiological processes of these complex diseases [5, 6]. Understanding the physiological basis for differences in the neurochemical response between healthy and diseased states is crucial for optimising clinical treatments.

MRS uses similar technology to magnetic resonance imaging (MRI), however the MRS signal is directly proportional to the concentration of metabolites within the specified region-of-interest (voxel), which is contrast to indirect MRI measures such as blood-oxygen-level-dependent (BOLD) imaging. Nonetheless, the physiological basis of the measured MRS neurotransmitter signal remains unclear [2, 3]. Focusing on Glu initially, the most widely-accepted explanation is that an increased MRS-Glu signal reflects increased metabolic turnover [7]. However, it remains controversial as to whether these processes are able to account for the large changes observed in the MRS signal (>5%) response to stimulation in short time scales of seconds to minutes [2, 3]. For example, Jelen *et al* [2] argue that changes in Glu of 12% or more cannot be explained by synthesis/degradation, since calculations based on known metabolic rates could not produce the same magnitude of change in such a short time. A major source of neuronal Glu is via flux through the Glu-Gln cycle, which has been shown in animals to exhibit rate increases in response to neural stimulation and so can reasonably be expected to do the same in humans [8]. Upon exocytosis into the cleft, Glu is quickly taken up by nearby astrocytes, converted to Gln (a non-neuroactive species) which is returned to the neuron where it is converted back to Glu and packaged into vesicles [9]. Supported by observations of varying glutamate MRS visibility in different compartments [10, 11] and on the echo-time dependence of fMRS signal changes, Jelen *et al.* proposed that fMRS detects ‘… compartmental shifts (in glutamate) due to neural activation, as it moves from pre-synaptic vesicles to more visible synaptic, extracellular and astrocytic pools’ [2]. Glu will naturally cycle through these compartments as it passes through the stages of release, reuptake and vesicular repackaging in response to neural firing. It has been estimated that up to 30% of Glu is invisible to MRS at any one time due to it being tightly bound to macromolecular structures in the vesicles causing a faster T2 relaxation rate [10, 11]. In the case of the inhibitory transmitter GABA, the process is less complex in that released GABA is returned to the postsynaptic cell largely via reuptake mechanisms where it is repackaged into vesicles [12]. Vesicle refilling has been shown to occur in an activity-dependent manner [13]. Therefore, it could be that neural activation drives a shift of Glu (or GABA) between states that are more or less visible to MRS via the recycling processes described above. Here, we test this hypothesis using a mathematical model. The model allows us to link cellular level mechanisms of neural activity to neurotransmitter dynamics with their systems-level observations in the MRS signal.

In this paper, we develop a microscopic-level model of neural activity based on the Hodgkin-Huxley formalism [14]. We explicitly incorporate a multi-compartment description of the movement of neurotransmitter between pools based on work by Tsodysks and Markram [15]. This multi-compartment model of NT dynamics assumes a fixed absolute quantity of NT that cycles between three ‘pools’: effective, inactive or readily releasable in an activity-dependent manner. We interpret these pools to correspond, respectively, to NT found instantaneously in the cleft upon cell activation or in the extracellular space (ECS); NT undergoing recycling or other biological processes (the cytosolic compartment); and NT packaged in vesicles ready for release. Essentially, the model accounts for the simplified dynamics of the release-reuptake-repackaging cycles described above. We obtain a macroscopic systems-level approximation of the neurotransmitter activity by deriving the meanfield approximation of the system dynamics under the Laplace approximation [16, 17]. Under the Laplace approximation, we formulate a description of the average response of a population of neurons to stimulation and the associated population-level changes in NT concentrations where the population density assumes a Gaussian form. We use current stimulation of varying polarity and magnitude to manipulate the mean membrane potential and firing rate of the model to make predictions about how the cells microscopic activity translates to changes in the fMRS measurement under the assumption that the vesicular pool of NT does not contribute to the MRS signal. In doing so, we provide a mechanistic account for the change in observed NTs concentrations during cognitive fMRS studies.

## 2 Methods and Materials

Initially, we model a cortical region as a canonical local network composed of two densely interconnected populations of spiking excitatory and inhibitory neurons. The neurons are coupled through excitatory *α*-amino-3-hydroxy-5-methyl-4-isoxazolepropionic acid (AMPA) and inhibitory (GABA) synapses. Glu acts on AMPA receptors to excite the cell and depolarise it, whereas GABA acts on GABA receptors to depolarise and inhibit it. A key component of the spiking model is that we account for the dynamics of both GABA and Glu during synaptic transmission. This allows us to study how the dynamics of the NT themselves vary in response to stimulation and how the MRS signal in turn would be affected.

Our aim is to relate the NT activity in the model to MRS-derived measurements of GABA and Glu which are acquired over large voxels in the brain containing billions of neurons. Using a network model comprised of individual neurons, this would entail simulating huge numbers of coupled nonlinear differential equations. Rather, we derive a mean-field reduction of the spiking network model. This reduction describes the evolution of the ensemble of neurons representing a local population, in terms of the dynamics of the average states of the ensemble. We follow Marreiros *et al.* to approximate the average ensemble behaviour under the Laplace approximation [16, 17].

### 2.1 Microscopic Model Dynamics

#### 2.1.1 Extended Hodgkin-Huxley model

As previously stated, the MRS signal is acquired over a voxel containing billions of neurons, we go on to define a network model of interconnected excitatory and inhibitory cells to enable us to compare the model output with the MRS data. Experimentally, it has been found that neocortical pyramidal cells exhibit dense reciprocal connectivity within a local region as well as strong connections with pyramidal cells in different regions via AMPA and N-methyl-D-aspartate (NMDA) receptor-mediated transmission [18–24]. Inhibitory cells receive excitatory input from pyramidal cells mainly via AMPA receptor-mediated transmission [25]. Similarly, GABAergic synapses have been found to mediate inhibitory connections on to pyramidal cells and inhibitory basket cells [26, 27]. Therefore, we base the model structure on these principles. We define a 2 populations of cells; an excitatory population and an inhibitory population coupled via AMPA, and GABA-mediated connections.

A single neuron is modelled using an extended form of the classical Hodgkin-Huxley (HH) neuronal model [14]. The temporal evolution of the neuronal membrane potential *V* (*t*) is driven by incoming signals from connected excitatory and inhibitory neurons, as well as external inputs. The equations describing the membrane potential for a single HH neuron *k* are given by (adapted from [28–30]):

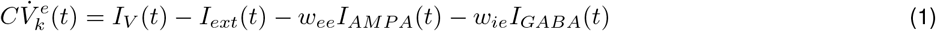

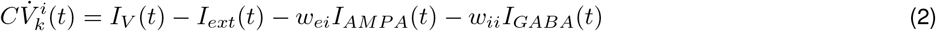

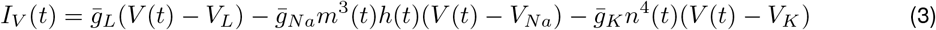

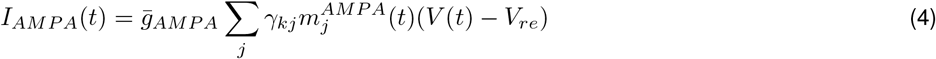

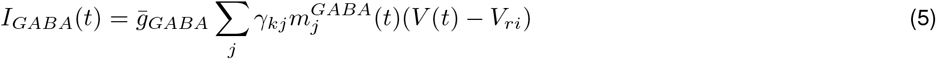

In Equations (1) to (5), (*V* ^*e*^) and (*V* ^*i*^) are the membrane potentials of the excitatory and inhibitory cells respectively; *C* is the specific capacitance of the membrane; *I*_*ext*_ is the external input current; *I_V_* encompasses the voltage-dependent leak, sodium and potassium currents respectively, which are responsible for the action potential generation; *g*_*L*_, *g*_*Na*_, *g*_*K*_ are the mean leak, sodium and potassium conductances and *V*_*L*_, *V*_*Na*_, *V*_*K*_ are the associated reversal potentials. The voltage-mediated gating variables *m*,*n*, and *h* represent the fraction of open channels of each current type (see Supplementary Information for a full list of equations).

We consider GABA_*A*_ receptors only as in, for example, [31] given that the majority of inhibitory postsynaptic potentials are mediated by GABA_*A*_ receptors rather than GABA_*B*_ receptors, which modulate slower inhibitory effects [29]. GABA_*A*_ receptors are also highly sensitive to GABA and can be fully saturated by GABA released from a single vesicle; whereas GABA_*B*_ require stronger stimuli to respond [29, 32, 33]. Henceforth, we omit the subscript for GABA_*A*_ and refer simply to GABA receptors.

The synaptic input currents are given by the AMPA (*I*_*AMPA*_), and GABA-receptor (*I*_*GABA*_) mediated synaptic input currents which will have either an excitatory or inhibitory effect on the cell. The respective synaptic conductances are *g*_*AMPA*_, and *g*_*GABA*_, and *V*_*re*_ and *V*_*ri*_ are the excitatory and inhibitory reversal potentials, respectively. A binary connectivity matrix *γ* represents a map of connections between individual neurons.

Recurrent connections within and between the two cell-type populations mediated via AMPA and GABA synapses have the weights *w*_*ee*_, *w*_*ii*_, *w*_*ei*_, and *w*_*ie*_, these are all fixed (see Table 1 for a list of parameter values).

**Table 1.** lists the parameter values used in the simulations unless stated otherwise.

| Paramater | Values | Reference |
| --- | --- | --- |
| $C$ | 0.01 F/m<sup>2</sup> | [28] |
| $g_l$ | 3 S/m<sup>2</sup> | [28] |
| $\bar{g}_K^e, \bar{g}_K^i$ | 0.006 S/cm<sup>2</sup>, 0.002 S/cm<sup>2</sup> | [28] |
| $\bar{g}_{Na}^e, \bar{g}_{Na}^i$ | 0.056 S/cm<sup>2</sup>, 0.01 S/cm<sup>2</sup> | [28] |
| $tv^e, tv^i$ | -56 mV, -68 mV | [28] |
| $V_l^e, V_l^i$ | -70 mV, -56 mV | [28] |
| $V_{Na}, V_K$ | 50 mV, -90 mV | [28] |
| $\bar{g}_a, \bar{g}_g$ | 25 nS, 10 nS | [29,31] (quantal conductances) |
| $\alpha_a, \beta_a$ | 1.1 s<sup>-1</sup>M<sup>-1</sup>, 180 s<sup>-1</sup> | [28,50] |
| $\alpha_g, \beta_g$ | 5 ms<sup>-1</sup>M<sup>-1</sup>, 166 s<sup>-1</sup> | [28,50] |
| $V_a, V_g$ | 0 mV, -80 mV | [28,29,50] |
| $w_{ee}, w_{ei}, w_{ie}$ | 0.1 | - |
| $w_{ii}$ | 0 | - |
| $U$ | 0.01 | [51] |
| $A$ | 10 mM | - |
| $\tau_{inact}$ | 3 ms | [15,51] |
| $\tau_{rec}$ | 1800 ms | (see Results) |
| $V_{max}, V_{tr}, \sigma_V$ | 1, 0 mV, 5 mV | - |

The synaptic gating channels for AMPA, and GABA mediated channels are given by the following generic equation:

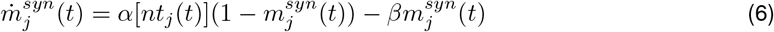

Where [*nt*] represents the concentration of neurotransmitter (GABA or Glu) acting instantaneously in the presynaptic cleft and *α* and *β* are rate constants specified in Table 1.

In standard conductance-based models, the synaptic currents are a function of the firing of presynaptic neurons which are in turn functions of the membrane potentials. Closing this loop allows for the membrane potential to be determined self-consistently. However, this type of modelling disregards the dynamics of neurotransmitter cycling. In this case, for the membrane potential to be computed self-consistently, the dynamics of the neurotransmitters need to be expressed as function of the pre-synaptic firing. In the next section, we describe the equations governing neurotransmitter cycling activity.

#### 2.1.2 Neurotransmitter Cycling

Upon activation, excitatory neurons release Glu into the postsynaptic cleft which rapidly binds with postsynaptic receptor sites and then is very quickly taken up by surrounding astrocytes. Within the astrocytes Glu is amidated to Gln by glutamine synthetase, and released into the extracellular space. Gln, a non-neuroactive species, is then taken up by the presynaptic neuron where it is converted back into Glu via phosphate activated glutaminase (PAG) and repackaged into vesicles. In addition to this, cytosolic Glu concentrations are also synthesised by the anaplerotic pathway via the tricarboxylic acid cycle (TCA) cycle.

In the case of inhibitory cells, released GABA is primarily taken up at the presynaptic terminal ready to be repackaged into vesicles. Synaptic GABA is also taken up into astrocytes where it is metabolised via the TCA cycle to succinate and then to Glu and finally Gln. In a similar way to the glutamatergic synapse, Gln is then released into the extracellular space and transported back to the presynaptic cell where it is converted back to Glu via PAG and finally to GABA via glutamic acid decarboxylase (GAD) after which it is packaged into vesicles ready for re-release. In addition to these two processes, cytosolic GABA is also synthesised by the anaplerotic pathway via TCA cycle as in the glutamatergic cell.

In this paper, we do not account for absolute changes in transmitter concentration due to metabolic flux in response to stimulation, as we assume that at short time-scales, these are unlikely to have a considerable effect on the MRS signal. Instead, we focus on movement of transmitters between cellular environments that are more or less visible to MRS to ascertain whether this shift in concentration is enough to explain the magnitude of change that we observe in the MRS signal over short timescales.

In order to account for the above mechanisms surrounding neurotransmitter recycling in the model, we make the following assumptions:

1. Neurotransmitter is released into the cleft in an activity-dependent manner and is rapidly removed so that it can no longer act on the postsynaptic cell.
2. Once removed from the cleft, neurotransmitter is assimilated into the cytosolic neurotransmitter concentration. We note here that for Glu, the situation is more complex as it is recycled via astrocytic conversion to Gln. Accounting for this extra step is an area for further work but we note here that in many MRS studies the combined measure Glx (which includes the Glu and Gln peaks) is used to approximate the Glu concentration and in these cases the model predictions are particularly relevant. Additionally, we note that Glu is present in approximately 4 times the concentration of Gln in the brain (see [34]).
3. Cytosolic neurotransmitter is then repackaged into the vesicles; the temporal evolution of this process depends on the level of cytosolic transmitter and levels of neural activity [13].

We employ the kinetic model of Tsodysks and Markram [15] which characterises the fraction of each NT as being in 1 of 3 states: *R* (the fraction of releasable transmitter in the bouton - the vesicular pool), *E* (the fraction of effective transmitter in the cleft) and *N* (the remaining proportion of ”inactive” transmitter which for our purposes we assume to be cytosolic neurotransmitter awaiting repackaging into vesicles), see Fig. 1. We adapt the model slightly by adding the term *N*_*base*_ which provides an upper bound to the proportion of NT that can be packaged into the vesicles. The equations describing the temporal evolution of NT in each compartment (where NT ∈ {Glu, GABA}) for cell *j* are given by:

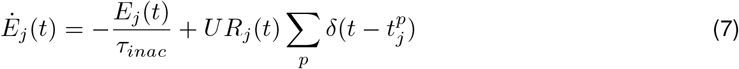

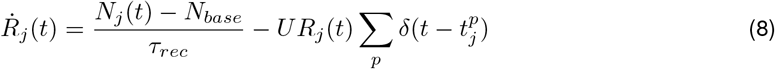

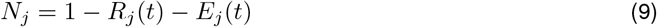

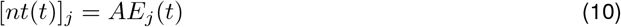

With each presynaptic spike, a fraction *U* of available neurotransmitter is activated (released) which then inactivates (reuptake) with time constant *τ*_*inact*_. The transmitter then undergoes recycling and repackaging into the vesicles at a rate *τ*_*rec*_.

**Figure 1:**
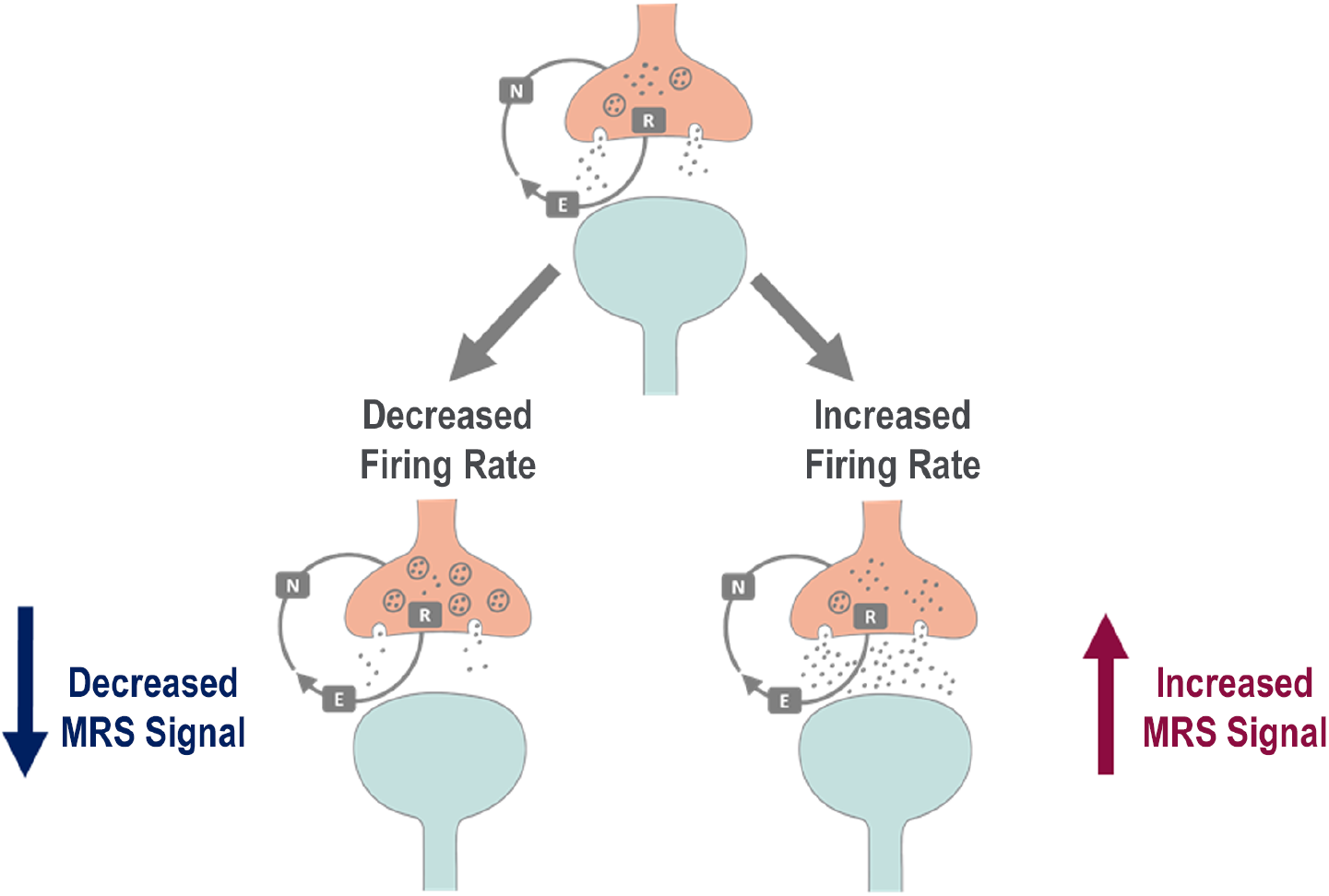
Neurotransmitter cycling. Schematic diagram showing the simplified and generalised NT cycling process as per the kinetic model of Tsodyks and Markram [15]. The presynaptic cell is shown in pink and the postsynaptic in blue. Steady state (top): the presynaptic cell fires an action potential releasing NT (GABA or Glu) into the cleft. NT is characterised as belonging to one of three states: in the vesicular pool, *R*, NT is packaged in the vesicles ready to be released into the cleft upon activation; NT in the ECS, *E*, has been released into the cleft and is acting at receptor sites on the postsynaptic cell or is in the ECS; and, NT in the cytosolic pool, *N*, is awaiting repackaging into the vesicles. Decreased firing (left): if the firing rate is reduced, then NT will start to accumulate in the vesicles due to the mismatch between the rate of exocytosis and the rate of repackaging into the vesicles. A shift of NT from the cytosol to the vesicles will result in a decreased MRS signal as we assume the vesicular compartment is largely invisible to MRS. Increased firing (left): if the firing rate increases, then NT from the vesicles will be used more quickly. Upon exocytosis it will transfer rapidly from the cleft to the cytosol and start to accumulate there. A shift of NT from the vesicles to the ECS and the cytosol will result in an increased MRS signal under the current hypothesis.

The input to cell *k* is calculated via a train of pulses (delta functions) that represents individual spikes arriving from pre-synaptic neuron *j* at times *p*. Equations (7) to (9) describe the proportion of the total *nt* in each of the pools which are multiplied by an absolute concentration, *A*, to convert the proportion of NT to a concentration in the cleft into a concentration measured in mM, denoted [*nt*]. Therefore, the concentration of neurotransmitter acting on the postsynaptic cell *k* (via eq. (6)) from presynaptic cell *j* is given by *AE*_*j*_.

As there is a considerable mismatch between the rate of exocytosis and the rate of recycling into the vesicles ready for release, a natural consequence of these equations is that during increased stimulation there is a shift of transmitters from a relatively MR-invisible compartment (the vesicle) to a fully visible compartment in the cytosol of astrocytes and neurons resulting in an increased MRS signal for that metabolite (see Fig. 1). The opposite effect occurs when firing is reduced.

### 2.2 Macroscopic Model Dynamics

#### 2.2.1 Mean-Field Model

A typical MRS signal is acquired from a large voxel of brain tissue (usually in the region of 27 cm^3^) meaning that it is a measure of activity at the macroscale. Our goal in this paper is to describe the evolution of metabolite dynamics in terms of what is measured by the MRS signal and therefore the model must also account for activity at this spatial scale. So far, the model equations account for the activity of a single cell; to model the behaviour of a network of billions of connected cells using these equations would be computationally expensive. Rather than taking this route, we follow Marreiros *et al.* and derive a mean-field model of macroscopic activity of a network under the Laplace approximation [16, 17]. The Laplace approximation is commonly used in statistical physics as it allows us to summarise the density dynamics of an ensemble of neurons using the method of moments [35, 36] where the population density assumes a fixed Gaussian form. The mean-field model (MFM) expresses the time-evolution of the mean activity of each population of neurons (excitatory and inhibitory) within the local spiking network described previously by averaging over the activity of the all the cells within each population. This is achieved by expressing the state variables by their mean values and replacing the sum over spikes found in eqs. (7) to (8) with an averaged value (firing rate) calculated using a sigmoidal input-output function [37]. The key point to note is that under the mean field assumption, the states of all neurons in a population are affected by the averaged state of all neurons in another population. It is not possible for a single neuron in a population to directly affect another individual neuron in the same or another population. In the MFM, we consider only the first statistical moment of the distribution, the mean, while discounting higher order moments thus equating the system to a neural mass model [17]. In the model, the excitatory and inhibitory populations that form a local population are both self-connected and interconnected, see Fig. 2.

**Figure 2:**
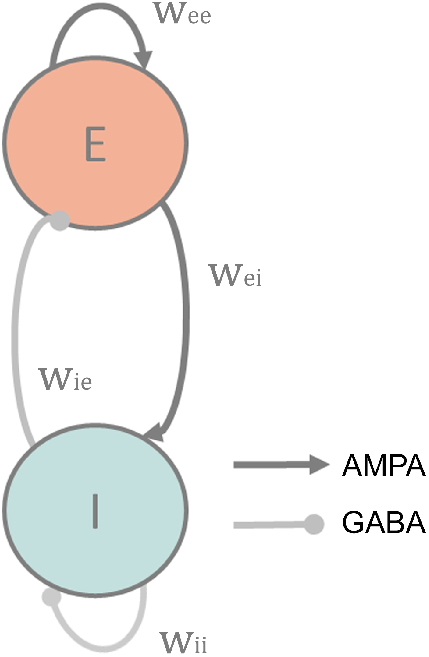
Canonical microcircuit of the mean-field model. Each mass, intended to approximately represent a standard MRS voxel, consists of an excitatory (E) and inhibitory (I) population that are both self and interconnected via the weights *w*_*ee*_, *w*_*ii*_, *w*_*ei*_, *w*_*ie*_. All connection strengths are fixed. Inhibitory connections are mediated via GABA receptors and excitatory connections via AMPA receptors.

In the spiking neuron model previously described, the input to cell *k* is in the form of a Glu (excitatory) or GABA (inhibitory) concentration found instantaneously in the cleft. For each conductance, *h*, this is calculated by summing over the spikes from a connected neuron. We can approximate the incoming spikes, for each *h*, as a Heaviside function on (*V − V*_*tr*_), where *V*_*tr*_ is a threshold that determines the proportion of cells firing at a point in time. Using mean-field theory, the input is a function of the population density of the source activity and under the Laplace approximation this is the Gaussian cumulative density which can be approximated by [17, 38, 39].

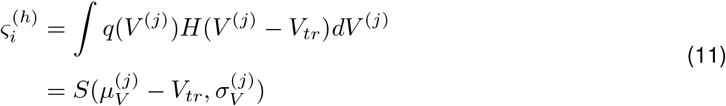

Where *S* is the sigmoidal function describing the cumulative density of the membrane’s depolarisation (or its deviation from resting levels), under Gaussian assumptions for that distribution with mean 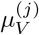 and variance 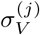, and 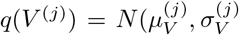. Under the Laplace approximation this function is simply the Gaussian cumulative density and can be approximated by [38–40]:

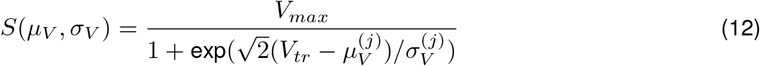

where *V*_*max*_ relates to the maximal membrane potential upon activation of the presynaptic cell.

Under mean-field assumptions, each neuron ‘senses’ the state of all neurons in each population it is connected to. Therefore we define the input to each neuron as a function of the mean presynaptic population activity by replacing the sum over spikes in eqs. (7) to (8) with an averaged rate corresponding to the population density of neurons of the same type from population *q*.

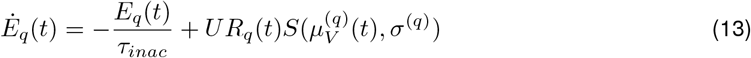

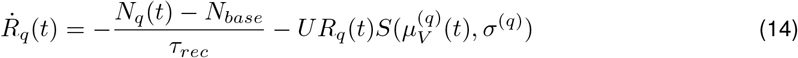

To estimate the network statistics, we approximate the dynamical equations for the statistical moments of the network’s state variables. First, we define the vector *x*_*p*_ of the model variables for local population *p* as 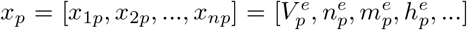 for a local population of neurons *p*. We express the system *p* of stochastic differential equations *f* (*x*) defined as eqs. (1) to (8) in terms of the first-order moments (the mean) of the distribution of the states, represented by the vector *μ*_*p*_.

We define

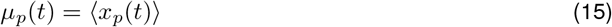

where angular brackets denote averages.

State variables are expressed as deviations from the means

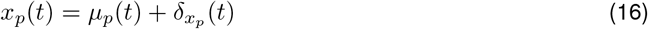

Performing a first-order Taylor expansion of *f* (*x*) around the mean *x* = *μ* we get (i.e. ignoring all second order and higher terms):

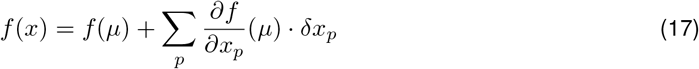

Averaging over realisations and noting that:

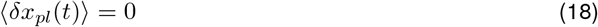

We obtain the motion equations for the means of the state variables for a local area *p*:

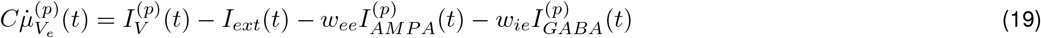

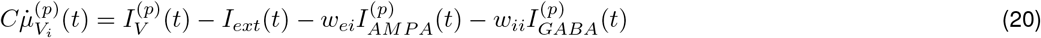

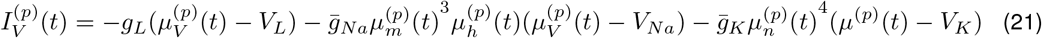

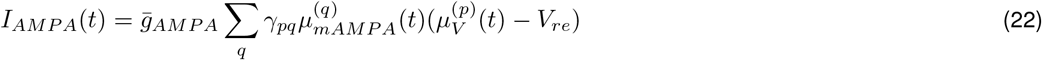

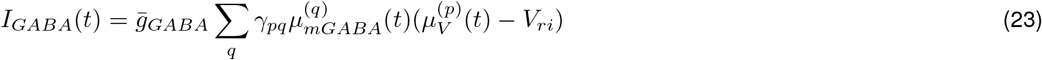

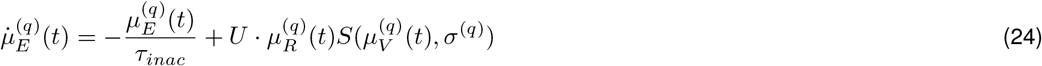

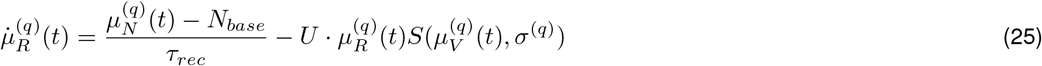

Where the superscript *nt* is omitted from *μ*_*E*_ and *μ*_*R*_, *nt* ∈ {Glu, GABA} and with *μ*_*n*_, *μ*_*m*_, *μ*_*h*_, and *μ*_*syn*_ defined similarly. We also note that in the present study, all connections are assumed to be instantaneous and therefore conduction delays between distant cortical areas are neglected.

#### 2.2.2 Driving the Network

Understanding how stimulation interacts with ongoing brain dynamics is important to optimising therapeutic and rehabilitation strategies [41]. To test the model, we added an exogenous input modelled as an input current that could be manipulated in terms of its direction and magnitude. This allows the model output to be interpreted in relation to both sensory and electrical stimulation types.

More formally, following [42] we added the term 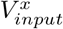(where *x ∈ {e, i}*) as an input current to each population of neurons which had either a hyperpolarising (analogous to cathodal transcranial direct current stimulation (tDCS)), or depolarising (analogous to sensory stimulation or anodal tDCS) effect depending on its direction. The direct effect of this on the MFM is that firing rates increase or decrease for a given ensemble of neurons which is consistent with experimental data [43, 44].

To test the effects of the input current, we applied the stimulus to both the excitatory and inhibitory populations following [45] who recently showed that extending a tDCS-style input to inhibitory interneurons as well as excitatory pyramidal cells allowed them to simulate evoked potentials that were much closer to physiological data. Here we used 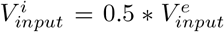. We note that the effects of electrical stimulation depend on a number of factors such as current field and cell orientation [46–48], and acknowledge that that this is an area for future work.

#### 2.2.3 Observation Model

MRS is the only method able to analyse the chemical composition of tissue in vivo. Using similar technology to MRI, MRS can detect chemicals and metabolites in the human brain existing in concentrations in the millimolar concentration range (mM). Individual metabolites can be differentiated by their characteristic pattern of chemical shifts, which reflect the local magnetic field experienced by each proton in a given molecule according to the local electronic and chemical environment, giving rise to a ’signature’ of resonant frequencies for a given molecule. This phenomenon gives rise to an MRS spectrum, see Fig. 3, which is formally defined as the Fourier transformed time-series containing the superimposed free induction decays from all the metabolites present in the sample. Peaks in the spectrum relate to protons bound to each molecular structure. Certain metabolites may be split between two or more peaks (such as GABA), and the summed area underneath these peaks is proportional to the concentration of the metabolite within the sample. Absolute quantification of the metabolites is possible, but complex, and therefore results are often expressed as a ratio to a reference metabolite such as N-acetylaspartate (NAA) or tCr (total creatine, the sum of creatine plus phosphocreatine).

**Figure 3:**
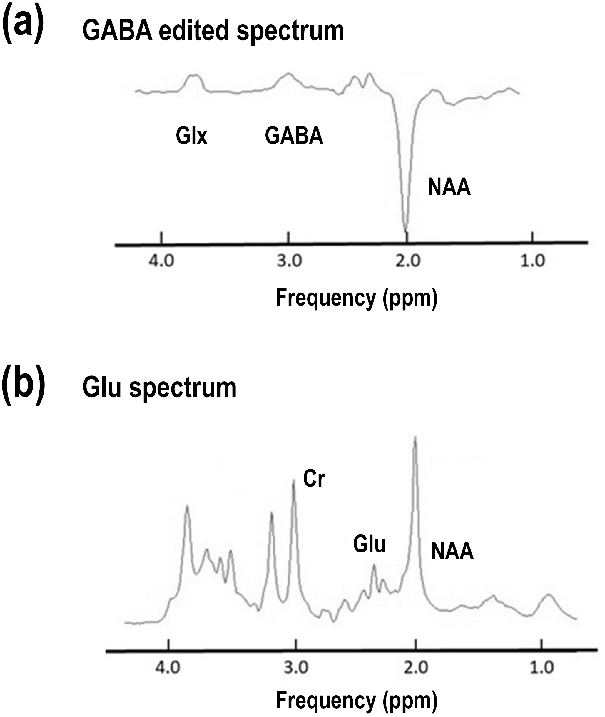
Example of Neurotransmitter Measurement by MRS. Example spectra are shown for the detection of (a) GABA, and (b) Glu in the human brain at a field strength of 3 Tesla. GABA is detected using a spectroscopic editing sequence. The edited spectrum, containing only NAA, GABA and Glx is shown. Spectra were acquired on a Philips 3 Tesla scanner at the Wellcome Trust / NIHR Clinical Research Facility, University of Manchester and Manchester University NHS Foundation Trust. Spectra were fitted using jMRUI [49]

To relate the model output to a standard fMRS experiment, we assume that only the ECS and cytosolic compartments contribute to the MRS signal. Therefore, we can relate the sum of the NT in the cytosolic and ECS pools directly to the concentration of NT in our region of interest. As the cytosolic pool is so much larger than the ECS pool then in practice, changes in the MRS signal are driven almost exclusively by changes in the size of this pool. In order to convert the proportions of NT within each of the 3 pools to a concentration then we must multiply by an average concentration for the voxel of the metabolite of interest measured in mM, *A*. Finally, when presenting the temporal trace of the NT dynamics in Fig. 6, the activity is presented as being averaged over a sliding window of 1 s which is comparable to the time resolution of the MRS signal (usually in the order of 1 to 5 seconds). All simulations were performed in Matlab (The Mathworks Inc., MATLAB ver. R2019b). The model was solved numerically using the forward Euler integration method with a time step of 0.01 ms, similar to [30].

#### 2.2.4 Parameter values

All parameters have been set to values within their physiologically measurable ranges, where such values exist.

## 3 Simulations and Results

In the present study we investigated the effect of a current based stimulation on neural activity and, in turn, on GABA and Glu dynamics. For this, we used the mean-field model described previously composed of 2 interconnected local networks, one excitatory and one inhibitory interconnected via AMPA and GABA receptor mediated synapses.

### 3.1 Effect of firing rate and amplitude of firing on NT cycling dynamics

Initially, we tested the effect of increasing firing rates on the NT cycling dynamics. We varied the average firing rate (modelled as a series of Dirac functions) between 0 and 100 spikes s^−1^ and solved the differential equations governing NT cycling rates (Equations (13) to (14)) for 100 s. *U*_*e*/*i*_ was set to 0.7, as in [15].

In Fig. 4 the percentages of NT found in the *R* pool (vesicular, left), the *E* (ECS, middle), and the *N* pool (cytosolic, right) were averaged over the duration of the simulation and are presented as a function of firing rate. Additionally, the amplitude of the input signal was varied which can be considered as a local measure of neural synchronisation in a MFM [52].

**Figure 4:**
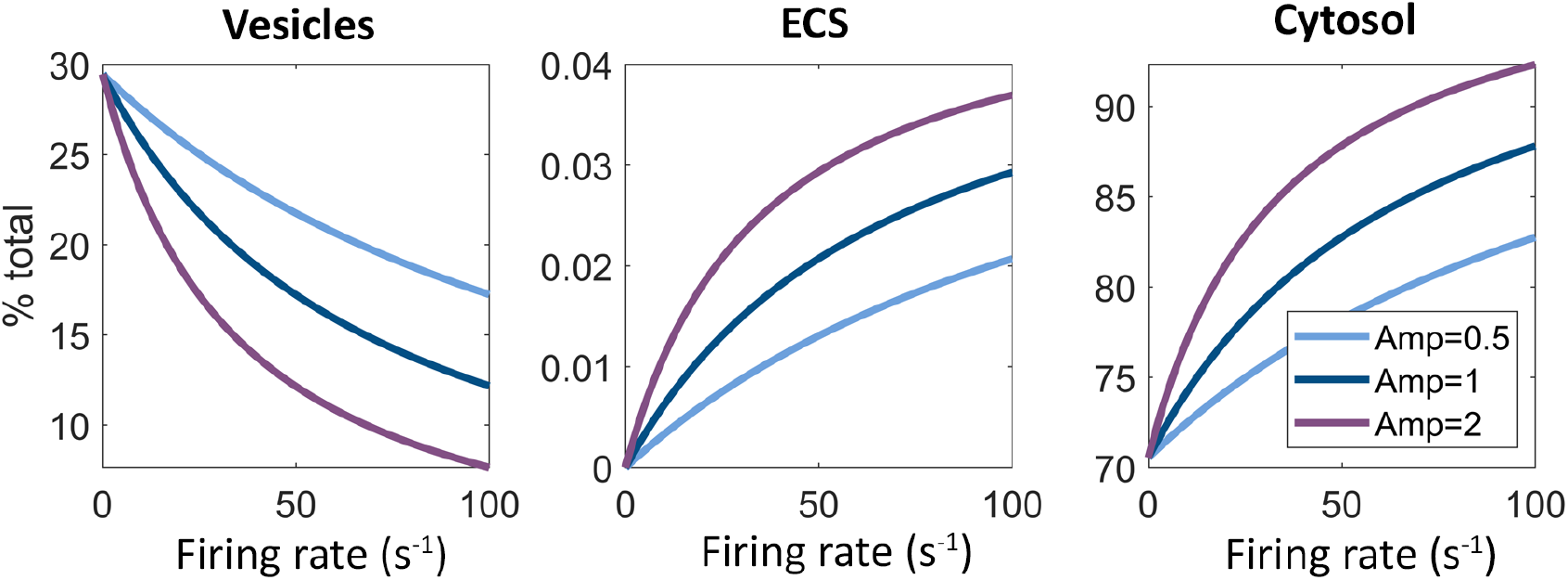
Effect of firing rate and amplitude of firing on NT cycling dynamics. The average proportion of neurotransmitter (averaged over 100 s of simulation) found in the vesicular pool (left), the ECS (middle), and the cytosolic pool (right) is presented as a function of firing rate and neural synchrony (amplitude). As the firing rate increases, the NT concentration shifts from the vesicles to the ECS and cytosolic compartments. If vesicular NT is not visible to MRS then this would be reflected as an increase in the MRS signal. Additionally, the amplitude of the input signal was varied between 0.5 (light blue), 1 (dark blue) and 2 (purple); we found that increasing the amplitude enhanced the shift of NT from the vesicular to the ECS and cytosolic pools for the same firing rate whereas reducing the amplitude inhibited this effect.

**Figure 5:**
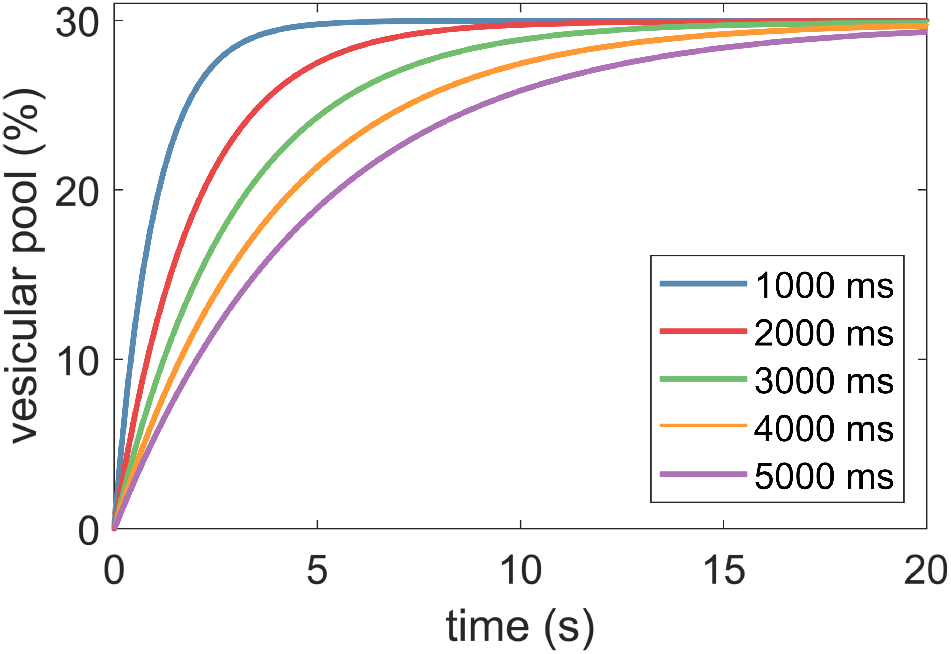
Effect of *τ*_*rec*_ on vesicle refilling times. The effect of *τ*_*rec*_ on the refilling time of the vesicular pool. Here *τ*_*rec*_ is varied between 500 and 5000 ms and the time course of the vesicular pool to reach its steady state value (from empty) is shown. We fix the value of *τ*_*rec*_ to be 1800 ms for the following simulations which results in refilling of the pool in approximately 10 seconds in line with [53] using a firing rate of 10 Hz.

When firing rates are at zero, the vesicular component is at its maximum (30% of the total) and as firing rates increase, this number is reduced. Variations in amplitude either enhance or inhibit this effect. We find that the reduction in vesicular NT is accompanied by an increased proportion of NT in the ECS (as a result of increased firing) or in the cytosol. The accumulation of NT in the cytosol is due to the mismatch between the rate that vesicles are depleted and the rate at which they are refilled [53]. We observe that, in general, an increase in firing (or amplitude of firing) leads to a shift of NT from the vesicular pool to the ECS and cytosolic pools, whereas a decrease in firing (or amplitude) drives a shift of NT into the vesicles as predicted (also see Fig. 1).

### 3.2 Effect of *τ*_*rec*_ on vesicular refilling times

Next, we measured the effect of *τ*_*rec*_ on the refilling time of the vesicular pool in order to ensure that the model behaviour is physiologically plausible. *τ*_*rec*_ governs the rate at which NT is recovered from the cytosolic pool and packaged into vesicles ready for release. Here *τ*_*rec*_ is varied between 1000 and 5000 ms and the time course for the vesicular pool to reach its maximum steady state value of 30% of the total cortical Glu concentration (starting from fully depleted vesicles) is shown. We fix the value of *tau*_*rec*_ to be 1800 ms for all simulations that follow as this value allows refilling of the vesicular pool in approximately 10 seconds which is in line with experimental data [53] using a firing rate of 10 Hz. We note that that the time taken to reduce this pool to a new lower steady-state value is in the order of 5 seconds which is also in line with experimental data [53], see Fig. 6. This result was achieved in the same way as the previous section, by solving the differential equations governing NT cycling rates (Equations (13) to (14)).

**Figure 6:**
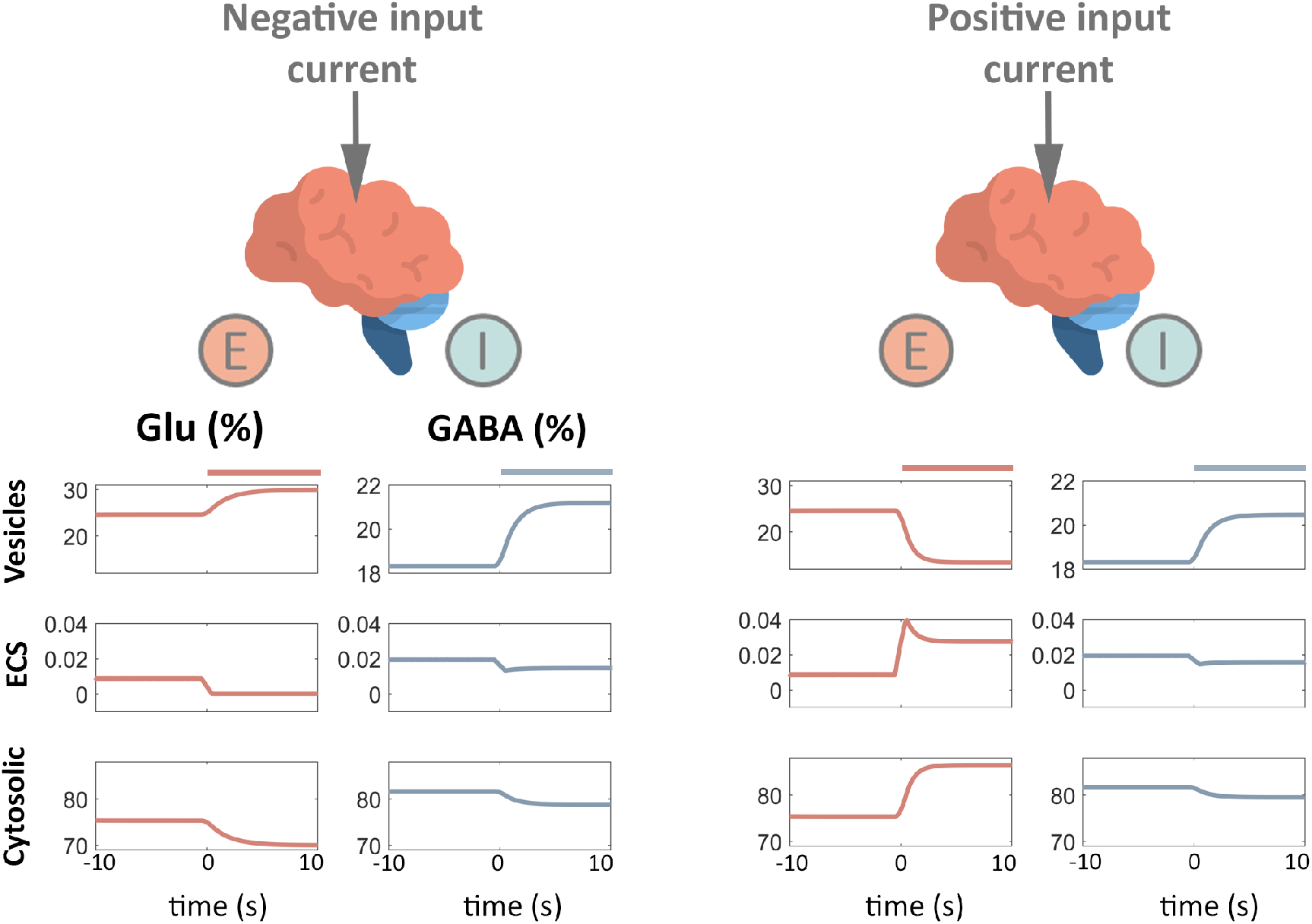
Effect of polarity-specific current stimulation on NT concentrations in each pool. We assess the effect of applying an input current on the NT dynamics within each pool and describe the expected effect on the resulting MRS signal assuming that the vesicular compartment is MR-invisible meaning that the cytosolic and ECS compartments determine the MRS signal. Initially, we apply an excitatory input (left panels). The network is allowed to run for 20 seconds so as to reach steady-state and then the excitatory stimulus 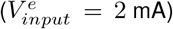 was applied for the remaining duration of the simulation (a further 10 seconds, see coloured bars). In the case of the excitatory population, we observe a shift of Glu from the vesicles to the cytosolic pool indicating an increased Glu MRS signal. In the case of the inhibitory population, we observe the opposite effect in that GABA accumulates in the vesicles and is reduced in the cytosolic compartment. This would be reflected as a reduction in the GABA MRS signal in this condition. In the case of an applied inhibitory current input 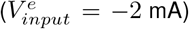, the percentage of Glu and GABA in the vesicular pools increases with a corresponding decrease in the cytosolic pool. The effect of this would be a reduction in both the Glu and GABA MRS signals. Note only 10 s of pre-stimulus activity is shown here.

### 3.3 Current stimulation and NT dynamics

We use an input current to manipulate the mean membrane potential and firing rate of the model to make predictions about how the MFM activity translates to changes in the fMRS measurement. The temporal trace of the NT dynamics in each of the 3 pools (vesicular, ECS, cytosolic) is given for the E-population (orange, left column) and the I-population (blue, right), see Fig. 6.

We allow the model to run for 20 s in order to reach a steady-state and then apply either a positive (excitatory, left panel) or negative (inhibitory, right panel) input current for the remaining duration of the simulation (see coloured bars) set at 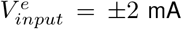. Note, data presented in Fig. 6 is for 10 s pre and post onset of stimulation.

For the excitatory input, in the case of the excitatory population, we observe a shift of Glu from the vesicles to the cytosolic pool. Under the assumption that NT in the vesicular pool does not contribute to the MRS signal, this would be reflected as an increased Glu-MRS signal. In the case of the inhibitory population, we observe the opposite effect in that GABA starts to accumulate in the vesicles and is similarly reduced in the cytosolic compartment. This change would be reflected as a reduction in the GABA MRS signal.

Next we applied an inhibitory input to the model after allowing the system to reach steady-state in the same way as before Fig. 6 (right panel). In this instance, we observe similar behaviour in both of the excitatory and inhibitory populations, specifically a shift of neurotransmitter from the cytosolic and ECS compartments to the vesicles. The effect of this in the imaging data would be a reduction in both the Glu and GABA MRS signals.

We used the model to systematically test the effects of current stimulation of varying direction and magnitude on GABA and Glu dynamics. We applied an input current that was varied between −5 to +5 mA to the MFM. Initially the model was allowed to run for 20 s to achieve a steady-state and then the input was applied for a further 20 s. In Fig. 7, the mean proportion of each of metabolites, GABA and Glu are presented for each of the pools, averaged over the duration of the simulation where the input current was applied. High-lighted with dashed lines is the case where 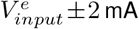 which corresponds to the diagrams shown in Fig. 6.

**Figure 7:**
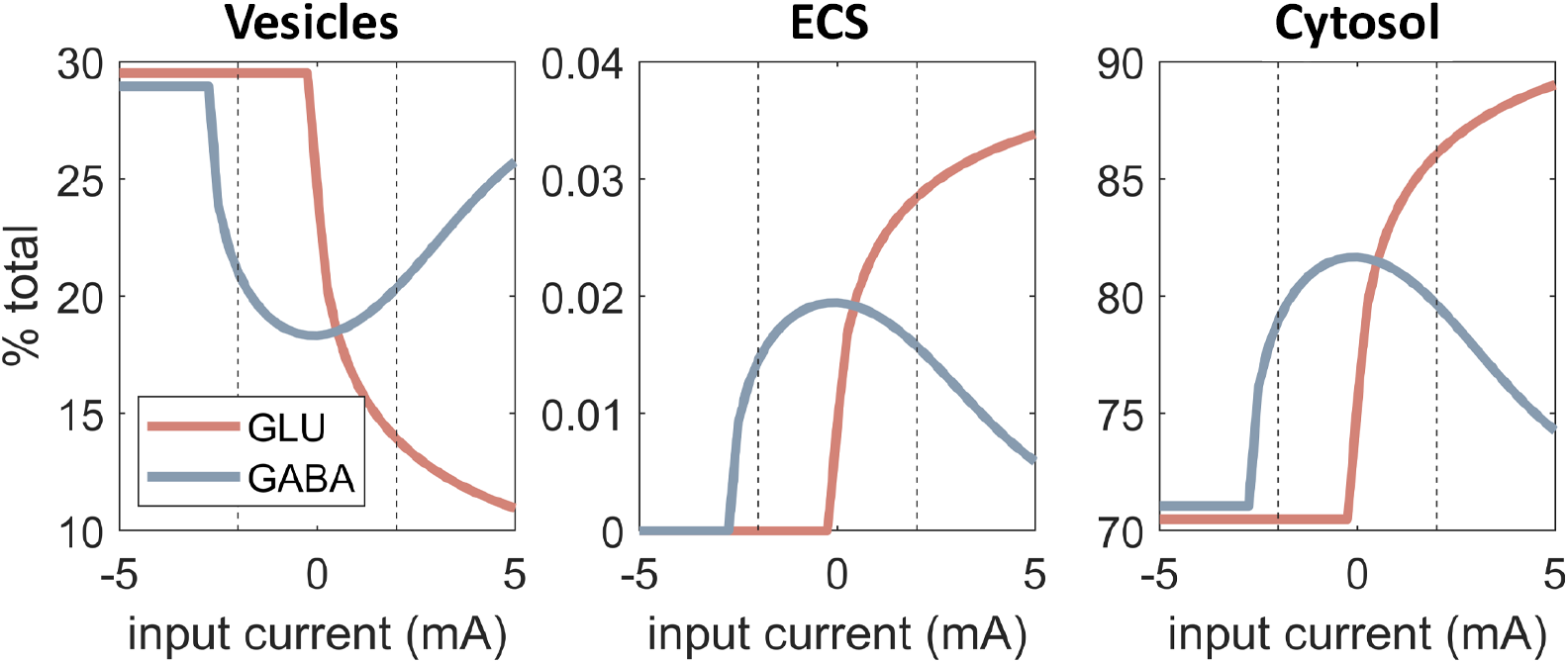
Effect of input current on NT dynamics. The proportion of Glu (orange) and GABA (blue) in each of the 3 pools averaged over the duration of the simulation is given for a range of positive and negative input currents. We find that for inhibitory inputs GABA and Glu are reduced in the ECS and cytosolic pools compared to the baseline condition 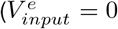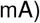. These changes would be reflected as a reduction in the MRS signal for both GABA and Glu. For excitatory inputs we observe a reduction in GABA and an increase in Glu in the ECS and cytosolic pools. This would be reflected as a decrease in the GABA MRS signal and an increased Glu MRS signal. Vertical dashed lines indicate the specific input values in Fig. 6 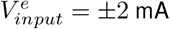.

For excitatory input currents we generally observe a shift of Glu from the vesicular to the cytosolic pools accompanied by a shift of GABA from the cytosolic to the vesicular pool. Under our assumption that NT in the vesicular pool does not contribute to the MRS signal, the model predicts that we would observe a decrease in the GABA signal and an increase in the Glu MRS signal in experimental scenarios using excitatory stimuli.

For inhibitory currents, we observe a reduction in GABA and Glu in the ECS and cytosolic pools accompanied by an increase of these metabolites in the vesicular pool compared to the baseline case 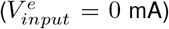. Therefore, the model predicts that we would observe a reduction in the MRS signal for both GABA and Glu in experimental scenarios using inhibitory stimuli.

For the range of inhibitory / excitatory current range applied here, the model predicts a change in Glu of between *±*10% compared to the baseline values within the cytosolic pool (as the ECS pool is so small it can practically be disregarded), and GABA reductions of up to 12% (we do not report a scenario where GABA concentrations increase).

Empirical studies using current stimulation report similar ranges; for example, in the visual cortex authors have found increases in Glu concentrations of up to 5% in response to photic stimulation measured over periods of seconds to minutes [54–56], and in response to flashing checkerboards [57]. Interestingly, Mekle *et al* observed a 5% reduction in GABA in response to a flashing checkerboard stimulus [58]. Similarly, a visual repetition suppression fMRS study reports increased Glu of 12% following novel stimulus presentation and decreases of 11% following repeated stimuli [59].

A number of fMRS studies focussed on pain report larger changes in the metabolite concentrations; for example, Cleve *et al* observe 21(15)% increase in Glx and a corresponding 15(12)% reduction in GABA measured in the anterior cingulate cortex (occipital cortex) in response to heat pain [60]. Whereas other authors have measured Glu increases of up to 20% in a range of brain regions over similar timescales of seconds to minutes [61, 62].

Finally, there are a number of studies looking at the motor cortex; Chen *et al* have reported increased Glx 8% and decreased GABA 12.5% in response to hand clenching [63]. Stagg *et al* show decreased GABA and Glx in response to cathodal (negative) current stimulation of 11 and 19% respectively and decreased GABA of 9% in response to anodal stimulation [64]. Floyer-Lea *et al* found 20 % decreases in GABA in response to motor learning [65], and Lea-Carnall *et al* observed a 30% decrease in GABA in response to repetitive tactile stimulation [66]. We note that this group of studies report changes in metabolites measured over longer time periods (10 - 46 mins) but include them here for comparison.

## 4 Discussion

In this study, we presented a mathematical model that bridges between the physiological changes at the micro-scale of the cell and synapse and fMRS measurements of the neurotransmitters GABA and Glu at the macroscale. The model was based on a standard Hodgkin-Huxley conductance-based formalism [14] which was extended to include AMPAR-mediated excitatory currents and GABAR-mediated inhibitory currents [29, 31] and tuned to have either the characteristics of an excitatory or inhibitory cell [28]. We further extended the model to account for the cellular-level NT dynamics developed by Tsodyks and Markram [15]. MRS, the technique used to measure Glu and GABA in vivo, acquires a signal at the level of the macroscale. To relate the model output to empirical observations, we derived the mean-field approximation to this system of equations using the Laplace approximation [16, 17]. This compartmentalised model of NT dynamics has been successfully incorporated into a number of mean-field models used to better understand phenomena such as working memory [67], EEG abnormalities [68], and selective attention [69], as well as theoretical studies of neural networks with dynamic synapses [51, 70]. We do not make a distinction between the cycling rates of GABA and Glu in the model. In reality, the molecular processes that underlie metabolism for these NTs are different but modelling these intricacies was beyond the scope of this work, which was intended to be a first exploration into understanding the effects of neurotransmitter cycling and the effect on the fMRS signal.

We stimulated the model using a range of positive (excitatory) and negative (inhibitory) inputs which had the effect of modulating the mean firing rate and mean membrane potential of the neural masses. This kind of stimulation was chosen to be analogous to standard sensory or electrical (tDCS) stimulation paradigms. We found that during excitatory stimulation, in the case of Glu, there was a general shift of NT from the vesicles to the cytosol which was explained by an increased firing rate for the excitatory population. This shift of Glu from the invisible vesicular compartment to the more visible ECS and cytosolic compartments would be reflected as an increased Glu MRS signal in response to excitatory stimulation. This is in line with numerous experimental papers that have shown increased Glu following excitatory sensory stimulation, see [3] for a review. In the case of GABA, we observed a reduction of NT in the ECS and cytosolic pools with a corresponding increase in the vesicular pool. This would be reflected as a decreased GABA MRS signal in response to an excitatory input. Excitatory input to the model can be thought of in the model as being analogous to sensory stimulation or anodal tDCS, and it has been suggested that anodal stimulation drives a reduction in the MRS GABA signal due to reduced activity of glutamic acid decarboxylase (GAD)-67 [64, 65], which is an enzyme involved in GABA synthesis that reduces with increased excitatory firing [71, 72]. The inclusion of these processes is beyond the scope of this work although we note that even without these details, the model predictions are in line with empirical observations.

During inhibitory stimulation (which can be thought of as being analogous to cathodal tDCS), we found reduced levels of both GABA and Glu in the extracellular and cytosolic pools, with a corresponding increase of both metabolites in the vesicular pool. Previously authors have observed reduced levels of GABA and Glu in response to cathodal stimulation [64]. It has been suggested that Glu is reduced as a direct consequence of decreased neuronal firing [64], as Glu/Gln cycling is tightly linked to glucose oxidation and therefore, neural activity [8, 73–75]. It has also been suggested that reductions in GABA found during cathodal stimulation are explained due to the maintenance of physiological balance between Glu, Gln and GABA (see [76–78]). In the model, this change is explained due to the reduction in inhibitory population firing activity although, in reality, this process is likely to be more complex. For example, in a recent study in the cat visual cortex, Zhao *et al.* found anodal (cathodal) tDCS enhanced (suppressed) the amplitude of visually evoked field potentials, these effects were not found with a sham condition [44]. The authors found anodal tDCS caused GABA to decrease with no change in Glu and cathodal tDCS to cause a reduction in GLU, but not GABA. Furthermore, that the mechanism driving these changes was that the polarity of the tDCS selectively suppressed the expression of GABA- and glutamate-synthesizing enzymes. Under our working assumption, that the NT pools in the cleft and the cytosol are MR visible with the vesicular pool being largely MR invisible, the model predicts a reduction in GABA- and Glu-MRS signal in response to a hyperpolarising current such as cathodal tDCS stimulation as seen in experimental studies [64, 79]. tDCS is thought to modulate the excitability of targeted neural populations by altering membrane potential in a polarity specific way [45]. Anodal tDCS has generally been found to increase cortical excitability and cathodal tDCS to reduce it with these effects validated via TMS motor evoked potential amplitudes [80].

In the model, we assume that there is no net increase in either Glu or GABA in the timescales shown here as it is assumed that flux through the Glu-Gln-GABA or TCA cycle would have minimal impact [2, 3]. However, we note that an alternative interpretation favours changes in metabolic flux as an explanation for stimulus-related Glu changes [81], based on calculations that the available pool of vesicular Glu is too small to account for the signal changes if this pool is released. We point out that this estimation is based on the assumption of 1 vesicle release per synapse at a single point in time [82]. In the model we account for sustained release of vesicular NT due to activation over many seconds driving the vesicular component closer to zero, or at least to a new lower steady-state value as firing rates increase. This has been found to be the case in as little as 5 seconds in animal studies using a preparation designed to mimic high frequency stimulation [53]. Although the model was initially designed to explain short term changes in physiology in response to stimulation, we suggest that longer term in-vivo measurements may also be explained by the same mechanisms presented here i.e. a mismatch between the rates of synthesis, repackaging, and exocytosis, regardless of the production pathways involved.

The key determinant of the shift of NT between the pools of varying MR visibility defined here, is the mismatch between the release and refilling rates of vesicles in response to an altered firing rate. Upon sustained stimulation, the number of vesicles released per second has been found to reduce until a lower steady-state is reached [53]. The release rate is predicted to decline according to the equation 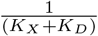 where *K*_*X*_ is the rate of exocytosis, and *K*_*D*_ is the vesicular refilling rate [53]. In the same paper, the authors estimated *K*_*X*_ to be almost seven times greater than *K*_*D*_. We chose *τ*_*rec*_ to allow a vesicle refilling rate of approx 10 seconds, as in [53]. Other authors have found this rate to be longer (23 seconds) [83]. Similarly, the model was tuned to reach a new steady-state in response to stimulation after approx 5 seconds, as observed in animal data [53]. Further work is required to ascertain the direct effect of these rates on NT dynamics.

Animal work has shown that the average sustainable release capacity for a central synapse is approximately two vesicles per *μ*m^3^ per second [84]. Assuming 5000 Glu molecules per vesicle (which is at the upper end of current estimations) [84], this equates to approximately 1 *μ*Mol g^−1^ min^−1^ which is equivalent to the 10% changes we observe in the model, assuming an average cortical Glu concentration of 10 mM [84, 85]. This rate would be further increased (allowing for even larger changes in shorter time scales) if not all of the cytosolic Glu were MRS-visible, as we have assumed here. This is higher than both the rate of flux through the Gln-Glu cycle [86] and the estimated rates of astrocytic uptake for Glu [87]. We suggest that in the short term it is not a requirement for Glu production to keep up with metabolic demand as there is a surplus of Glu in the cytosol which can be recycled into vesicles. Astrocytic uptake rates must also be considered as a limiting factor; however, it has been argued that Glu uptake is relevant to the “slow component of glutamate removal [from the cleft]…” [84, 88, 89] and that immobilisation by binding is a much faster method of inactivating Glu allowing some time for astrocytic uptake to take place. Therefore, in the short-term, these limiting factors may not apply.

The metabolic pathway involving each of the neurotransmitters modelled here is highly complex; this holds in the case of Glu particularly. For example, within the neuron, cytosolic Glu may be utilised for new amino-acid syntheses, entry into the TCA cycle, or conversion into glutathione or GABA amongst many other possibilities [9]. Here we make the simplified assumption that all cytosolic Glu in the model is available for repackaging into vesicles for neurotransmission.

In this paper we bridge the gap between spatial scales by developing a mean-field model of macro measurements in fMRS, based on the hypothesis that fMRS reflects a shift between pools. We have developed a mean-field model to link human in vivo quantification of task-related neurotransmitter changes via MRS to synaptic activity. The model allows the prediction of MRS-Glu and GABA as a result of neural activation. It gives a mechanistic explanation for the magnitude of the NT changes observed in fMRS measurements, thus provides a theoretical basis for the study of the role of these neurotransmitters in cognitive processing.

## 5 Acknowledgements

The authors would like to thank Prof. Peter Robinson for his careful reading and insightful comments on an earlier version of this manuscript.

## 6 Funding

CLC is funded by the Medical Research Council (MRC) Grant MR/PO14445/1. WeD acknowledges the financial support of ANID, Chile, projects FONDECYT 1201822, ANILLO ACT172121 and Basal FB0008. NTB is funded by EPSRC Grant EP/N006771/1. CJS holds a Sir Henry Dale Fellowship, funded by the Wellcome Trust and the Royal Society (102584/Z/13/B). The Wellcome Centre for Integrative Neuroimaging is supported by core funding from the Wellcome Trust (203139/Z/16/Z).

## Notes

### Competing Interest Statement

The authors have declared no competing interest.

