## Supplementary Information for "From synaptic activity to human in vivo quantification of neurotransmitter dynamics: a neural modelling approach"

### 1 Spiking model equations

Here we include the auxiliary equations for the extended Hodgkin-Huxley model described in the main paper.
The sodium and potassium voltage-gated currents respectively, which are responsible for the action potential generation are given by:

$$I_{Na}(t) = \bar{g}_{Na} m^3(t) h(t)(V(t) - V_{Na}) \tag{1}$$

$$I_K(t) = \bar{g}_K n^4(t)(V(t) - V_K) \tag{2}$$

where, $V(t)$ is the membrane potentials of the neurons (mV); $g_{Na}, g_K$ are the mean sodium and potassium conductances (mV) and $V_{Na}, V_K$ are the associated reversal potentials (mV). The voltage-mediated gating variables $m$,$n$, and $h$ represent the fraction of open channels of each current type and are given by:

$$\dot{m}(t) = \alpha_m(V(t))(1 - m(t)) - \beta_m(V(t))m(t) \tag{3}$$

$$\dot{h}(t) = \alpha_h(V(t))(1 - h(t)) - \beta_h(V(t))h(t) \tag{4}$$

$$\dot{n}(t) = \alpha_n(V(t))(1 - n(t)) - \beta_n(V(t))n(t) \tag{5}$$

With:

$$\alpha_m(V) = \frac{-0.32(V - tv - 13)}{\text{exp}[-(V - tv - 13)/4] - 1} \tag{6}$$

$$\beta_m(V) = \frac{0.28(V - tv - 40)}{\text{exp}[(V - tv - 40)/5] - 1} \tag{7}$$

$$\alpha_h(V) = 0.128\, \text{exp}[-(V - tv - 17)/18] \tag{8}$$

$$\beta_h(V) = \frac{4}{1 + \text{exp}[-(V - tv - 40)/5]} \tag{9}$$

$$\alpha_n(V) = \frac{-0.032(V - tv - 15)}{\text{exp}[-(V - tv - 15)/5] - 1} \tag{10}$$

$$\beta_n(V) = 0.5\, \text{exp}[-(V - tv - 10)/40] \tag{11}$$

All taken from [1].
